# Tissue- and Age-Specific Transcriptomic and Metabolomic Analysis Reveals Regulatory Mechanisms of Ginsenoside Biosynthesis in *Panax notoginseng*

**DOI:** 10.64898/2026.03.02.708933

**Authors:** Wei Li, Yiduo Wang, Fugang Wei, Gao Xiong, Zhongjian Chen, Lizhi Gao

**Author notes:** These authors contributed equally to this work (Wei Li, Yiduo Wang).

## Abstract

Ginsenosides are the primary bioactive compounds in *Panax notoginseng*, but the transcriptional mechanisms governing their tissue- and age-dependent accumulation remain elusive. Here, we integrated targeted metabolomics with transcriptome profiling across four tissues (root, stem, leaf, and flower) and three developmental stages (1–3 years) to investigate the spatiotemporal regulation of ginsenoside biosynthesis. We observed distinct tissue- and age-specific accumulation patterns: roots exhibited a progressive increase in total ginsenoside content during the second and third years, while flowers preferentially accumulated rare protopanaxadiol-type ginsenosides such as Rg3-2. Transcriptomic analysis revealed extensive differential gene expression across tissues and stages, particularly in roots during late development. Clustering and transcription factor (TF) enrichment analyses identified multiple tissue-associated regulatory modules. Four TFs—*AT3G12130*, *SPL9*, *MYB33*, and *SPL1*—emerged as core candidates based on coordinated expression, promoter motif enrichment, and functional annotation of predicted target genes. Motif analysis further linked these TFs to key biosynthetic genes involved in triterpenoid oxidation and glycosylation, including *CYP716A53v2* and *CYP716A47*. Together, these findings suggest that tissue-specific ginsenoside accumulation in *P. notoginseng* is associated with coordinated transcriptional regulation of biosynthetic enzymes. This study provides a transcriptomic framework for understanding spatial regulation in ginsenoside biosynthesis and identifies candidate regulators for future functional validation and metabolic engineering.

## Introduction

*Panax notoginseng* (Burk.) F.H. Chen is a perennial herbaceous plant of the Araliaceae family with a long history of medicinal use. It is recognized in the Chinese inventory of “medicinal and edible” plants and has been increasingly utilized in the health food industry [1]. Traditionally, *P. notoginseng* has been valued for promoting blood circulation, resolving stasis, relieving pain, and nourishing the blood. Modern pharmacological studies have further confirmed its antioxidative, anti-inflammatory, antihyperlipidemic, and immunomodulatory activities [2–4]. These health benefits have led to its widespread application in functional foods, dietary supplements, and natural therapeutics. The primary bioactive constituents of *P. notoginseng* are dammarane-type triterpenoid saponins, collectively referred to as notoginsenosides. These compounds are not only essential for the plant’s pharmacological effects but also serve as quality indicators, with their content and distribution varying significantly across tissues [5].

Notably, different tissues of *P. notoginseng* exhibit distinct bioactive profiles and physiological effects. For example, the roots are well recognized for cardiovascular protection, the leaves have been reported to alleviate anxiety and improve sleep [6], and the flowers have shown potential benefits in regulating blood pressure and reducing dizziness [7]. These differences suggest that organ-specific metabolic pathways contribute to the plant’s diverse functions. Therefore, identifying tissue-specific saponins and their biosynthetic regulators is essential for the targeted and safe application of *P. notoginseng* in health food products. In addition, ginsenoside accumulation is influenced by plant age. Typically, plants aged two to three years accumulate higher levels of active compounds and are preferred for consumption, underscoring the importance of investigating both spatial and temporal metabolic variations.

The biosynthesis of notoginsenosides follows the canonical triterpenoid secondary metabolic pathway, beginning with the production of isoprenoid precursors (IPP and DMAPP) via the mevalonate (MVA) or methylerythritol phosphate (MEP) pathway. These intermediates are then converted **to** squalene and cyclized by enzymes such as dammarenediol synthase (DDS) to form the triterpenoid backbone [8–10]. Subsequent modifications involve hydroxylation by cytochrome P450 monooxygenases (e.g., CYP716A family) and glycosylation by UDP-glycosyltransferases (UGTs), resulting in structurally diverse ginsenosides [11]. Based on structural features, these saponins are typically classified as protopanaxadiol (PPD) and protopanaxatriol (PPT) types, which vary in their tissue distribution, likely due to the differential expression and regulation of biosynthetic genes.

Recent studies have widely employed RNA sequencing (RNA-seq) and metabolomics to investigate the biosynthetic mechanisms of *P. notoginseng* saponins. Substantial differences in saponin composition have been observed across organs and developmental stages, with roots generally accumulating more saponins than aerial parts, particularly between the second and third years of growth [12–14]. Key biosynthetic genes, including farnesyl diphosphate synthase (*FPS*), squalene synthase (*SS*), dammarenediol synthase (*DS*), squalene epoxidase (*SE*), *UGTs*, and members of the *CYP716A* family, show strong tissue- and age-specific expression patterns [12,15,16]. For example, the floral bud stage has been identified as a critical phase for Rb3 accumulation, with significant upregulation of genes related to triterpene backbone biosynthesis and structural modifications [16]. Regarding functional gene validation, genes such as *DS* and *CYP716A47*-like have been functionally characterized through transgenic expression [17]. Moreover, high-quality full-length transcriptomic datasets obtained using single-molecule real-time (SMRT) sequencing have greatly improved gene annotation, revealing novel transcript variants and long non-coding RNAs involved in ginsenoside biosynthesis [15].

The genome of *P. notoginseng* has been sequenced, providing a valuable foundation for investigating the biosynthesis and metabolism of bioactive compounds, as well as for conducting genome-level functional studies [1]. Integrated transcriptomic and metabolomic approaches, including WGCNA and KEGG pathway enrichment analyses, have facilitated the identification of key candidate genes and regulatory networks controlling saponin accumulation [14,18–20]. These studies have established robust correlations between gene expression and metabolite accumulation across tissues and germplasms.

Despite these advancements, the complexity of *P. notoginseng*’s metabolic network and its multilayered spatiotemporal regulation require further systematic studies across developmental stages and tissues. To address this gap, the present study integrates transcriptomic and targeted metabolomic data from four tissues (root, stem, leaf, and flower) over three growth years (1, 2, and 3 years, respectively) to elucidate the spatial-temporal dynamics of saponin accumulation. Through expression pattern analysis, we identified candidate genes and regulators closely associated with tissue-and age-dependent saponin profiles. Using high-throughput sequencing, gene ontology enrichment, and promoter motif scanning, we further identified several transcription factors potentially involved in regulating key biosynthetic steps, such as oxidation and glycosylation. Our findings highlight root and flower tissues as major sites of saponin biosynthesis, with strong concordance between saponin types and transcriptional regulation. Overall, this study provides insights into regulatory mechanisms of ginsenoside biosynthesis in *P. notoginseng* and offers a molecular framework for future metabolic engineering and functional food development.

## Results

### Multi-organ transcriptomic and targeted metabolomic profiling of *P. notoginseng* across three years

To characterize tissue- and age-dependent regulation of ginsenoside biosynthesis in *Panax notoginseng*, we performed integrated transcriptomic and targeted metabolomic analyses across four tissues collected over three consecutive years (Fig. 1A). Seventeen representative ginsenosides were quantified in each tissue using targeted metabolomics (Fig. 1B), and RNA-seq was performed in parallel to assess genome-wide gene expression patterns associated with ginsenoside accumulation.

**Figure 1.**
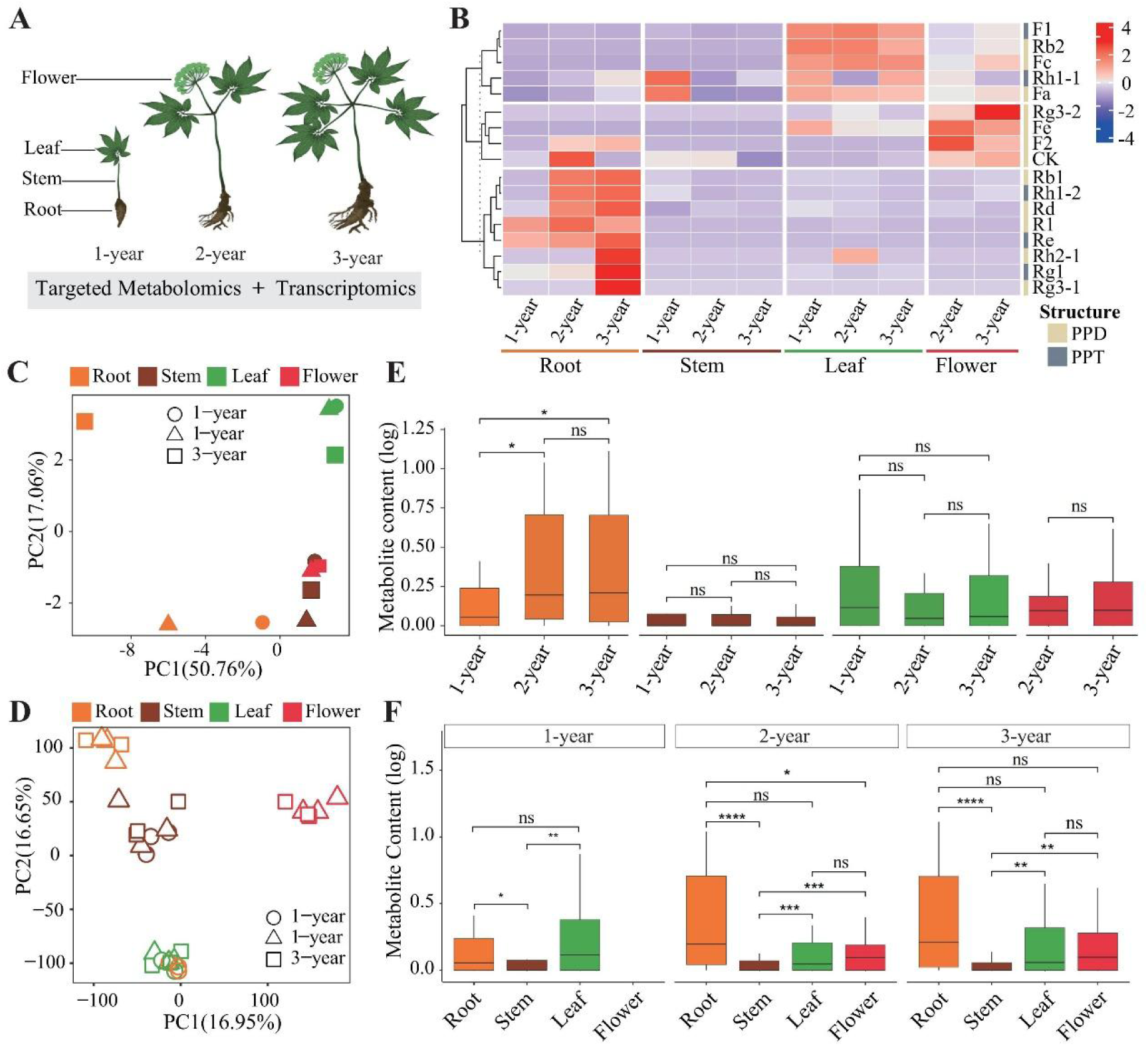
Experimental design and integrated metabolomic and transcriptomic overview of *P. notoginseng*. **(A)** Schematic overview of sample collection and experimental design across four tissues (root, stem, leaf, and flower) over three consecutive growth years. **(B)** Heatmap showing the relative contents of 17 targeted ginsenosides across tissues and years. Ginsenoside levels were normalized using row-wise Z-score transformation to facilitate comparative analysis. PPD, protopanaxadiol-type notoginsenosides; PPT, protopanaxatriol-type notoginsenosides. **(C)** Principal component analysis (PCA) of samples based on the contents of the 17 ginsenosides, illustrating metabolite-based separation among tissues and growth years. **(D)** PCA of transcriptomic profiles based on global gene expression levels, revealing transcriptional variation across tissues and developmental stages. **(E)** Comparison of ginsenoside contents across different growth years within the same tissue. **(F)** Comparison of ginsenoside contents among different tissues within the same growth year. Statistical significance was assessed using pairwise Wilcoxon rank-sum tests. Significance levels are denoted as * *P* < 0.05, ** *P* < 0.01, and *** *P* < 0.001; ns indicates no significant difference.

Principal component analysis (PCA) of both ginsenoside profiles and transcriptomic data showed that, in most tissues, tissue variation exceeded inter-annual variation. In contrast, root samples displayed a pronounced age-dependent separation, with first-year roots clearly distinct from second- and third-year roots, exceeding the variation observed among tissues (Fig. 1C, D).

Consistent with the PCA results, ginsenoside accumulation increased significantly with plant age only in root tissues (*p* < 0.05) (Fig. 1E). Within each year, ginsenoside contents differed significantly among tissues, with stems consistently showing the lowest levels, while roots were significantly higher than stems but generally comparable to leaves and flowers depending on developmental stage (Fig. 1F). Individual ginsenosides showed marked tissue- and age-specific accumulation patterns. Rh2-1, Rg1, and Rg3-1 were most abundant in roots of three-year-old plants, whereas flowers preferentially accumulated protopanaxadiol (PPD)-type ginsenosides, with Rg3-2 showing strong enrichment in third-year flower samples (Fig. 1B).

RNA-seq was performed with three biological replicates per sample. All libraries showed mapping rates above 90%, except for one replicate each from three-year roots and one-year leaves, which were excluded due to low quality (Table S1). Biological replicates were highly reproducible, with Pearson correlation coefficients ranging from 0.70 to 0.97 (Fig. S1). Across all samples, 30452 genes (83% of annotated genes) were expressed (CPM > 1) in at least one sample, with 74–85% of genes detected across tissues and years, indicating broadly conserved transcriptional activity across tissues and developmental stages (Fig. S2).

### Tissue- and year-specific differential expression patterns in *P. notoginseng*

To systematically characterize transcriptional variation across tissues and developmental stages in *P. notoginseng*, we performed differential expression analysis using DESeq2 (q < 0.05, |log₂ fold change| > 1). Differentially expressed genes (DEGs) were identified for year-to-year comparisons within each tissue and for inter-tissue comparisons within each year.

Year-to-year comparisons revealed marked tissue-dependent differences in transcriptional dynamics (Fig. 2A). Root tissue showed the most pronounced transcriptional variation, with 8,854 and 8,957 DEGs identified in the comparisons between the first and second years and between the first and third years, respectively. In contrast, only 323 DEGs were detected between the second and third years in roots. Stem tissues showed substantially fewer DEGs, with 580 and 729 DEGs identified in the first vs. second and first vs. third year comparisons, respectively, and 138 DEGs between the second and third years. Leaf and flower tissues exhibited intermediate levels of year-to-year transcriptional change. Overall, these results indicate that major transcriptional reprogramming occurs predominantly between the first year and later developmental stages, particularly in roots.

**Figure 2.**
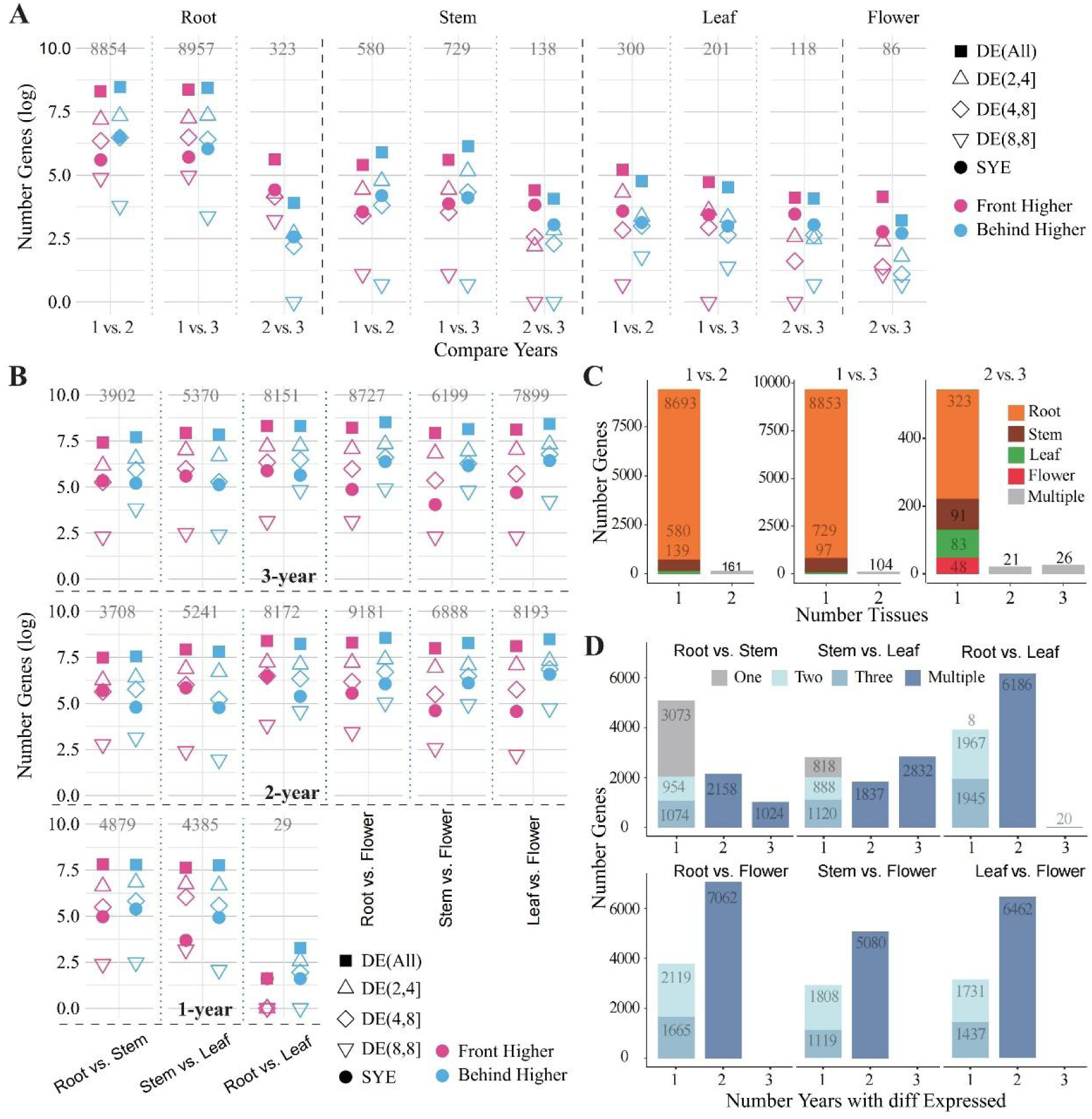
Developmental dynamics and conservation of gene expression patterns across tissues and years in *P. notoginseng*. **(A)** Identification of differentially expressed genes (DEGs) among tissues within the same growth year. Differential expression was determined using the same criteria applied throughout this study (q < 0.05 and |log₂ fold change| > 1). **(B)** Conservation of tissue-specific expression differences across growth years. The x-axis indicates the number of years in which a given gene exhibited significant tissue-specific differential expression. **(C)** Conservation of inter-annual expression differences across tissues. The x-axis indicates the number of tissues in which the same gene showed significant differential expression between years. **(D)** Year-to-year conservation of tissue-specific differential gene expression. Bar plots show the number of DEGs identified for each pairwise tissue comparison within the same year. The x-axis indicates the number of years (1, 2, or 3) in which each DEG was detected for a given tissue comparison, while the y-axis represents the total number of DEGs. Colors denote the specific year in which a DEG was detected when it occurred in only one year (One, Two, or Three), or Multiple when detected in two or more years. Each panel corresponds to a distinct tissue pair.

Differential expression analysis between tissues within the same year revealed a strong developmental effect on tissue-specific transcriptional divergence (Fig. 2B). In the first year, only 29 DEGs were identified between root and leaf tissues, indicating limited transcriptional differentiation at this stage. In contrast, in the second and third years, the number of DEGs between root and leaf increased dramatically to 8,172 and 8,151, respectively. Comparisons among other tissue pairs within the same year also yielded large numbers of DEGs, ranging from 3,902 (root vs. stem in the third year) to 9,181 (root vs. flower in the second year). These results demonstrate that transcriptional differences among tissues become increasingly pronounced as plants mature.

To evaluate the tissue specificity of year-dependent expression changes, we assessed the overlap of DEGs among tissues for each year-to-year comparison (Fig. 2C). Notably, all 323 DEGs identified between the second and third years in roots were unique to root tissue. In stems, among the 138 DEGs detected for the same comparison, 66% (91 genes) were stem-specific, while the remaining genes were shared with other tissues. This pattern indicates that late-stage year-to-year transcriptional changes are highly tissue-specific in roots, whereas stems show a greater degree of shared expression changes across tissues.

We further examined the extent to which tissue-specific transcriptional differences were conserved across years (Fig. 2D). Comparisons involving flower tissues showed relatively high conservation, with 65%, 63%, and 67% of DEGs between flower and root, flower and stem, and flower and leaf tissues, respectively, detected in at least two years. Similarly, 61% and 62% of DEGs identified in leaf vs. root and leaf vs. stem comparisons were conserved across years. In contrast, only 38% of DEGs between root and stem tissues were shared across multiple years, indicating greater temporal variability in transcriptional differences between these two tissues.

### Spatiotemporal expression clusters reveal tissue-specific modules associated with ginsenoside accumulation

To investigate the biological relevance of genes exhibiting tissue- and year-specific expression patterns in *P. notoginseng*, all expressed genes were grouped into 15 spatiotemporal expression clusters (C1–C15) based on their expression dynamics.

In roots, clusters C2–C5 showed pronounced upregulation during the second and third growth years, a pattern that closely paralleled the progressive accumulation of ginsenosides in this tissue (Fig. 3A; Fig. 1B). Functional enrichment analysis revealed that these clusters, particularly C4, were predominantly enriched in fundamental cellular processes related to gene expression, including ribonucleoprotein complex biogenesis, RNA metabolism and processing, ribosome biogenesis, and macromolecule metabolic processes (Fig. S3). These results suggest that enhanced transcriptional and translational capacity in roots during later developmental stages may provide a cellular basis for large-scale ginsenoside accumulation.

**Figure 3.**
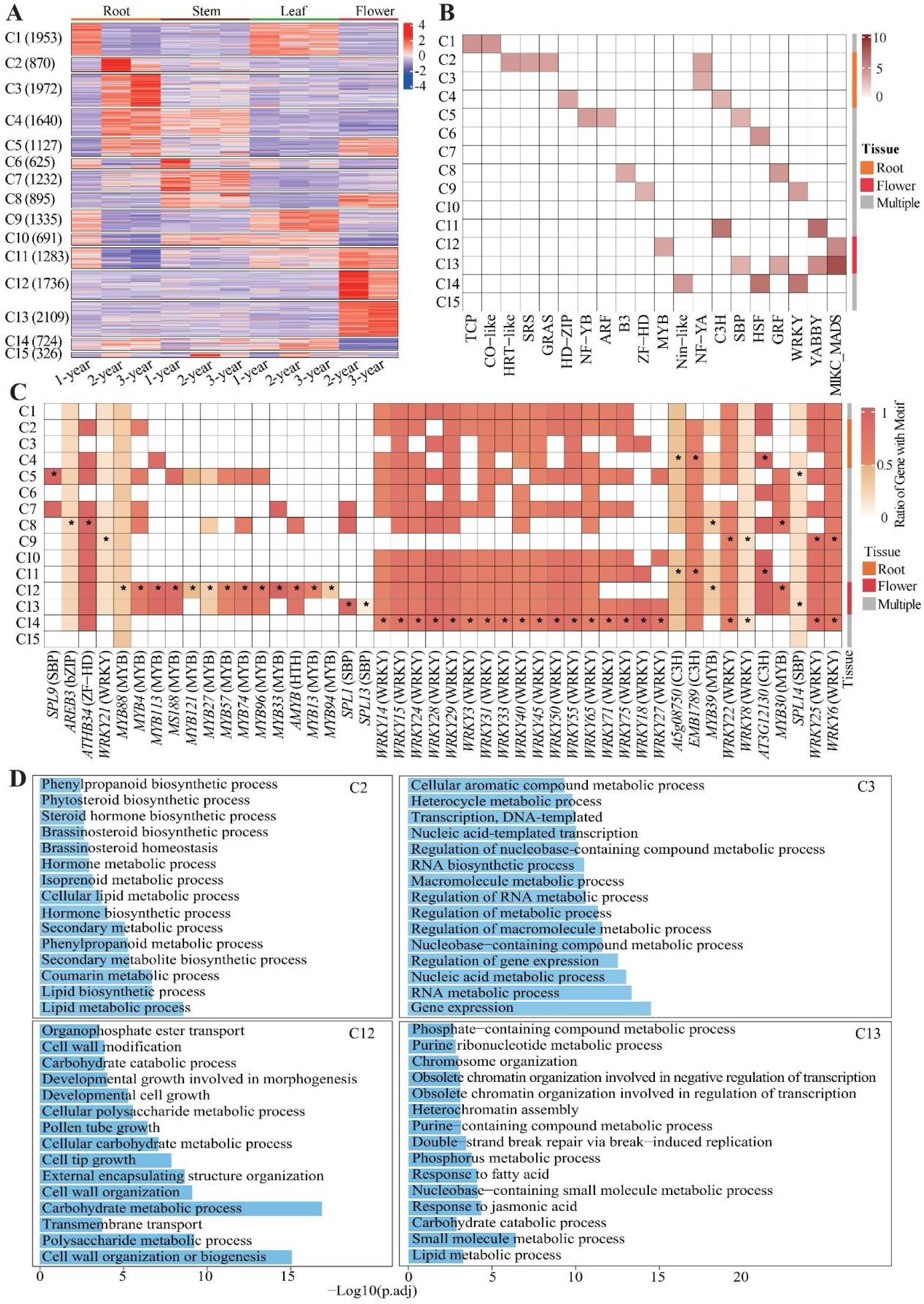
Dynamic patterns of gene expression and transcription factor enrichment analysis across all samples. **(A)** Expression dynamics of genes showing differential expression in at least one pairwise comparison across all samples. Expression values were Z-score normalized by row. Based on expression patterns, genes were clustered into 15 groups (C1 ∼ C15). Each column represents a sample, and each row represents a gene. **(B)** Enrichment of transcription factor (TF) families among C1 ∼ C15 clusters. TF genes were annotated from panel A, and enrichment was tested using a two-sided Fisher’s exact test. Red indicates significantly enriched TF families (q < 0.05). Each row represents a TF family, and each column represents a gene cluster. **(C)** Enrichment of known transcription factor binding motifs in the 2 kb upstream promoter regions of genes in each cluster. White indicates non-significant enrichment; colored blocks indicate significant enrichment (q < 0.05). Color intensity reflects the proportion of genes within the cluster containing the motif, with yellow and red representing clusters where >50% of genes contain the motif. Asterisks (*) indicate that the TF family corresponding to the motif is also significantly enriched in the same cluster (as shown in panel B), supporting direct transcriptional regulation. **(D)** Gene Ontology (GO) enrichment analysis of genes in clusters C2, C3, C12 and C13. Selected representative biological process terms are shown. The color scale represents the enrichment significance (-log10(q-value) or enrichment score).

In leaves, clusters C1 and C9 exhibited consistently high expression across all three growth years, in agreement with the relatively stable ginsenoside levels observed in leaf tissues (Fig. 3A; Fig. 1B). Functional enrichment indicated that C1 was mainly associated with photosynthesis-related processes, whereas C9 was enriched in RNA processing and organelle-related pathways (Fig. S3). These findings imply that maintaining ginsenoside homeostasis in leaves may depend on coordinated energy supply and transcriptional regulation. Notably, both C1 and C9 also showed relatively high expression in roots during the first growth year, suggesting that these modules may function broadly during early development before becoming more leaf-associated at later stages.

In flowers, clusters C5, C8, C11, C12, and C13 were strongly upregulated, with expression patterns consistent with year-dependent changes in ginsenoside accumulation in this tissue (Fig. 3A; Fig. 1B). Among them, C12 and C13 showed pronounced flower-specific expression. Functional enrichment analysis showed that C13 was mainly involved in chromatin organization, nucleotide and lipid metabolism, small-molecule metabolism, and responses to jasmonic acid, whereas C12 was significantly enriched in processes related to cell wall remodeling, carbohydrate and polysaccharide metabolism, transmembrane transport, and pollen tube and tip growth (Fig. 3D). These results suggest that ginsenoside biosynthesis, transport, and tissue-specific accumulation in flowers may be jointly regulated by transcriptional and structural cellular programs.

Overall, these findings indicate that leaf, root, and flower tissues in *P. notoginseng* form distinct yet coordinated expression modules aligned with ginsenoside biosynthesis, accumulation, and allocation, collectively reflecting a clear source–sink organization at the transcriptomic level.

### Enrichment of transcription factor families highlights tissue-specific regulatory mechanisms of ginsenoside biosynthesis

To further elucidate the regulatory mechanisms underlying the observed expression modules, transcription factor (TF)–encoding genes were annotated across all clusters (C1–C15), and TF family enrichment was systematically assessed (Fig. 3B). Given that transcriptional regulation depends not only on TF expression levels but also on their ability to recognize cis-regulatory elements, we additionally examined the enrichment of known TF binding motifs within the 2-kb upstream promoter regions of genes in each cluster (Fig. 3C).

The C3H TF family was significantly enriched in clusters C4 and C11, both of which showed sustained upregulation in roots and flowers during the second and third growth years. Motif analysis further revealed that binding sites of three C3H TFs (AT3G12130, EMB1789, and At5g08750) were highly prevalent in the promoter regions of genes in these clusters. Notably, AT3G12130 binding motifs were detected in 98.6% and 99.0% of promoters in clusters C4 and C11, respectively (Fig. S4A), suggesting that C3H TFs may directly regulate genes involved in ginsenoside-related processes in these tissues.

In cluster C12, MYB transcription factors were strongly enriched, particularly MYB33, MYB133, and MS188, whose corresponding binding motifs were detected in 98%, 87%, and 85% of C12 gene promoters, respectively (Fig. 3B; Fig. 3C; Fig. S4A). Genes in C12 exhibited flower-specific high expression (Fig. 3A), and flowers predominantly accumulated protopanaxadiol-type (PPD-type) ginsenosides, such as Rg3-2, Fe, and F2 (Fig. 1B). These observations suggest that MYB TFs may play a central role in flower-specific regulation of ginsenoside metabolism. The preferential enrichment of MYB binding motifs in flower-associated genes further supports a MYB-mediated tissue-specific transcriptional regulatory framework.

WRKY transcription factors were significantly enriched in cluster C14, whose genes were upregulated in roots, stems, and leaves. In addition, WRKY binding motifs were detected across multiple clusters, including C9, indicating that WRKY TFs may have broader regulatory roles across both vegetative and reproductive organs.

Interestingly, in several clusters, TF binding motifs were enriched even when the corresponding TFs were not transcriptionally upregulated, highlighting the context-dependent nature of transcriptional regulation and suggesting that different TF family members may regulate distinct target gene sets under specific spatial or developmental conditions.

Consistent with the tissue-specific composition of ginsenosides—Rb1, Rd, and Rg1 being predominant in roots, whereas Rg3-2, Fe, F2, and CK were enriched in flowers (Fig. 1B)—the observed TF enrichment patterns showed clear correspondence: C3H TFs were mainly associated with root-related clusters, while MYB TFs dominated flower-related clusters. Together, these results indicate that tissue-specific transcription factor networks orchestrate distinct ginsenoside biosynthetic programs, thereby shaping the spatial diversity of ginsenoside accumulation in *P. notoginseng*.

### Identification of core transcription factors associated with tissue-specific ginsenoside biosynthesis

To elucidate the transcriptional regulatory basis of ginsenoside biosynthesis, all expressed genes were classified into 15 expression clusters (C1–C15) according to their spatial and temporal expression patterns across different tissues and growth years (**Fig. 3A**). A transcription factor (TF) was defined as a core TF if it simultaneously satisfied the following criteria: (i) its binding motif was significantly enriched in the promoter regions of genes within the corresponding cluster; (ii) its TF family was significantly overrepresented among the genes in that cluster; and (iii) more than 90% of cluster genes contained the TF-specific binding motif within the 2 kb upstream of the transcription start site. Based on these criteria, six core TFs were identified, including *AT3G12130* (C3H family) in clusters C4 and C11, *SPL9* (SBP family) in C5, *ATHB34* (ZFHD family) in C8, *MYB33* (MYB family) in C12, and *SPL1* (SBP family) in C13 (Fig. S4A).

Targeted metabolomic profiling revealed pronounced accumulation of ginsenosides in root and flower tissues (Fig. 1B). Accordingly, we focused on four core TFs exhibiting tissue-specific upregulation, namely *AT3G12130* (*C3H*) and *SPL9* (*SBP*) in roots, and *MYB33* (*MYB*) and *SPL1* (*SBP*) in flowers. Gene Ontology (GO) annotation of the predicted target genes of these TFs showed significant enrichment in triterpenoid-related metabolic functions, particularly oxidoreductase and glycosyltransferase activities (Fig. S4B; Table S2).

Consistent with these functional annotations, genes encoding key enzymes in the ginsenoside biosynthetic pathway showed clear tissue-specific expression patterns that closely matched the accumulation profiles of different ginsenoside types (Fig. 4A). In root tissues, genes involved in protopanaxatriol (PPT)-type ginsenoside biosynthesis were preferentially upregulated. Notably, *CYP716A53v2* (*PPTS*) exhibited high expression levels during the second and third growth years (Fig. 4B), corresponding to the enrichment of PPT-type ginsenosides, including Rh1-2, Rg1, Re, and R1, in roots (Fig. 5A).

**Figure 4.**
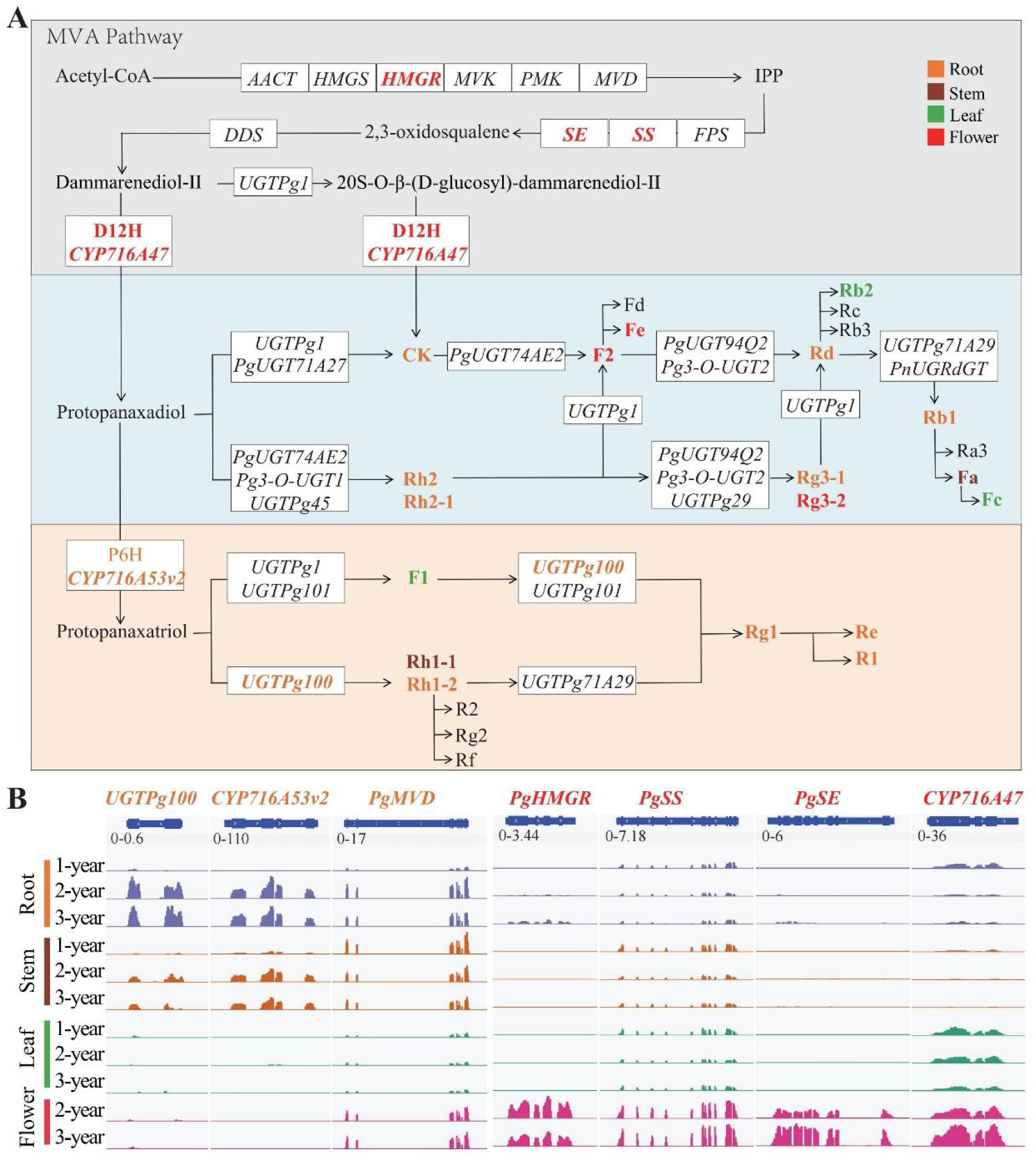
Expression patterns of key enzyme-encoding genes involved in ginsenoside biosynthesis in *P. notoginseng*. **(A)** Biosynthetic pathways of protopanaxadiol (PPD)-type and protopanaxatriol (PPT)-type ginsenosides. Enzymes are shown as colored rectangles, with colors indicating tissues (root, stem, leaf, and flower) in which the corresponding genes are significantly upregulated. **(B)** IGV visualization of representative enzyme-encoding genes involved in PPD- and PPT-type ginsenoside biosynthesis. RNA-seq read coverage is shown for root, stem, leaf, and flower tissues across three consecutive growth years. Colored tracks represent different tissues, and peak heights reflect relative transcript abundance of the same gene across tissues and years. Genes associated with PPT-type ginsenoside biosynthesis (e.g., *CYP716A53v2* [*PPTS*]) show predominant expression in roots, whereas genes involved in PPD-type biosynthesis (e.g., *CYP716A47*, also known as *PqPPDS*/*PqD12H*) are highly expressed in flowers.

**Figure 5.**
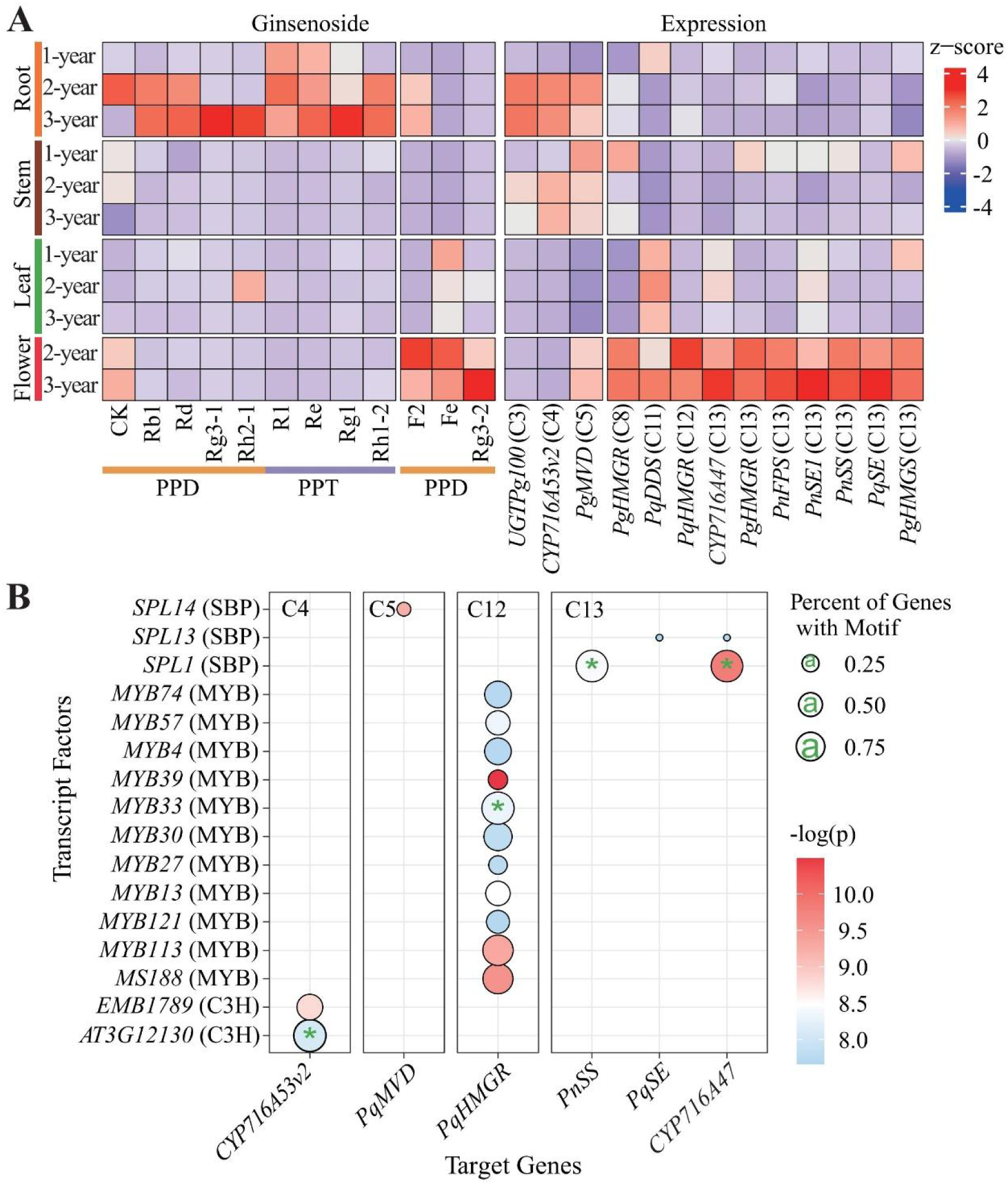
Coordinated expression of ginsenoside biosynthetic enzymes and associated transcription factors in *P. notoginseng*. **(A)** Integrated analysis of tissue-specific expression patterns of key ginsenoside biosynthetic enzyme–encoding genes across root, stem, leaf, and flower tissues over three growth years, together with the corresponding ginsenoside accumulation profiles. Heatmaps show normalized expression levels of representative enzymes involved in protopanaxadiol (PPD)-type and protopanaxatriol (PPT)-type ginsenoside biosynthesis in different tissues and years. **(B)** Enrichment of core transcription factor binding motifs in the promoter regions (2 kb upstream of the transcription start site) of key ginsenoside biosynthetic enzyme–encoding genes. Bar plots indicate the proportion of target genes containing each motif, and color intensity reflects motif enrichment significance. P-values were calculated using FIMO.

In contrast, genes associated with protopanaxadiol (PPD)-type ginsenoside biosynthesis showed higher expression levels in flower tissues. Among them, *CYP716A47* (*PPDS*), which catalyzes the conversion of dammarenediol-II to PPD-type intermediates, was markedly upregulated in flowers (Fig. 4B), in agreement with the high accumulation of PPD-type ginsenosides such as F2, Fe, and Rg3-2 in this tissue (Fig. 5A). These results indicate a strong concordance between tissue-specific ginsenoside accumulation and the expression patterns of key biosynthetic enzyme-encoding genes.

Further promoter motif analysis revealed potential transcriptional associations between core TFs and key biosynthetic enzymes. A C3H binding motif corresponding to *AT3G12130* was detected within the 2 kb upstream promoter region of *CYP716A53v2*, whereas an SBP binding motif corresponding to *SPL1* was identified in the promoter region of *CYP716A47* (Fig. 5B). The distribution of these motifs was consistent with the tissue-specific expression patterns of both the TFs and their associated biosynthetic genes.

Collectively, these results demonstrate that tissue-specific accumulation of distinct ginsenoside types in *P. notoginseng* is accompanied by coordinated expression of core transcription factors and key biosynthetic enzymes. The identified core TFs and their promoter motif associations provide important transcriptomic evidence for the spatial regulation of ginsenoside biosynthesis.

## Discussion

This study integrated targeted metabolomic and transcriptomic analyses of root, stem, leaf, and flower tissues of *Panax notoginseng* across three developmental stages (one-, two-, and three-year-old plants), providing a comprehensive view of the spatiotemporal dynamics of ginsenoside accumulation. By combining metabolite profiling with gene expression and regulatory analyses, our results deepen the understanding of the biosynthetic regulation of key bioactive constituents in *P. notoginseng* and provide a molecular basis for its rational utilization in functional food development and industrial applications.

Metabolomic profiling revealed a wide spectrum of ginsenosides, including major protopanaxadiol (PPD)-type compounds (Rb1, Rb2, and Rd), protopanaxatriol (PPT)-type compounds (R1, Rg1, and Re), and multiple rare ginsenosides such as CK, F2, Fa, Fc, Fe, Rg3-1, Rg3-2, Rh2-1, F1, Rh1-1, and Rh1-2. These compounds are recognized for their diverse pharmacological activities, including lipid regulation, antioxidant, anti-inflammatory, anti-fatigue, and immune-modulatory effects, underscoring their high value for health product development[21–24]. Rare ginsenosides such as CK, Rg3, and Rh2 have attracted particular interest due to their low natural abundance. Recent studies have shown that CK can activate brown adipose tissue, enhance thermogenesis, and modulate the gut microbiome, thereby contributing to obesity prevention and improved insulin sensitivity [25,26], further supporting the potential of these compounds in functional food and nutraceutical development.

Clear tissue-specific and age-dependent patterns of ginsenoside accumulation were observed. Roots of two- and three-year-old plants accumulated significantly higher total ginsenoside contents than stems, leaves, and flowers (p < 0.05), consistent with previous reports identifying roots as the primary storage organ for ginsenosides[12,18]. The higher ginsenoside levels detected in three-year-old roots also align with traditional harvesting practices, supporting the empirical view that three-year-old *P. notoginseng* roots represent optimal medicinal quality. Moreover, individual ginsenosides exhibited distinct organ-specific distributions. For example, Rg3-1 and Rh2-1 were highly enriched in three-year-old roots and have been associated with cardiovascular protective effects[27,28], whereas flowers accumulated high levels of Rg3-2, particularly in three-year-old plants, consistent with their traditional use in relieving dizziness and regulating blood pressure. In addition, CK was mainly detected in two-year-old roots and flowers. Given that CK is a gut microbial metabolite derived from Rb1 and Rb2[29], its tissue-specific distribution, together with that of its precursors, provides valuable information for the targeted development of functional products. Collectively, these results indicate that different organs of *P. notoginseng* have distinct ginsenoside profiles, reflecting functional specialization and offering opportunities for tissue-specific utilization.

Consistent with metabolomic patterns, transcriptomic analyses revealed extensive tissue- and age-dependent differential gene expression. Previous studies have shown that ginsenoside content in *P. notoginseng* roots increases with plant age and serves as an important index for quality evaluation[13]. Our data confirmed that roots of two- and three-year-old plants exhibited both higher ginsenoside contents and a large number of specifically upregulated genes, suggesting a strong association between transcriptional regulation and metabolite accumulation. GO enrichment analysis indicated that these differentially expressed genes were mainly involved in secondary metabolism, terpenoid backbone biosynthesis, lipid metabolism, and phenylpropanoid metabolism. The phenylpropanoid pathway, mediated by oxidases, ligases, reductases, and transferases, is known to play a central role in regulating plant secondary metabolism [30–32]. The enrichment of these pathways suggests coordinated regulation among multiple metabolic networks contributing to ginsenoside biosynthesis in roots.

Phytohormone-related pathways may further modulate this process. In this study, genes associated with brassinosteroid (BR) biosynthesis were significantly upregulated in roots. Although BRs have not been directly implicated in ginsenoside biosynthesis, they are known to interact with other hormones such as methyl jasmonate (MeJA) and salicylic acid (SA). For example, BR-overaccumulating mutants in Arabidopsis exhibit suppressed SA signaling and increased susceptibility to pathogens [33]. Both MeJA and SA have been reported to enhance ginsenoside accumulation by inducing the expression of biosynthetic enzyme genes[34,35]. Therefore, BRs may indirectly influence ginsenoside metabolism through crosstalk with MeJA or SA signaling pathways, a hypothesis that warrants further investigation.

At the pathway level, triterpenoid saponin biosynthesis initiates from the mevalonate (MVA) pathway, with key structural genes such as *FPS*, *SS*, *SE*, and *DS* playing critical roles. In agreement with previous reports[13], *SS* and *SE* showed higher expression in aerial tissues, particularly flowers, whereas ginsenosides predominantly accumulated in roots. This discrepancy suggests that active biosynthesis may occur in aerial organs, followed by transport of ginsenosides or their intermediates to belowground tissues for storage. In addition, two key cytochrome P450 enzymes, D12H and P6H, displayed contrasting tissue-specific expression patterns. D12H was preferentially expressed in flowers, consistent with the dominance of PPD-type ginsenosides, whereas P6H was highly expressed in roots, supporting the accumulation of both PPD- and PPT-type ginsenosides in this tissue. These findings highlight the importance of spatial regulation of P450 enzymes in determining ginsenoside composition among organs.

To further explore transcriptional regulation, we identified four core transcription factors—*AT3G12130*, *SPL9*, *MYB33*, and *SPL1*—that exhibited pronounced tissue-specific expression patterns and motif enrichment in the promoters of ginsenoside biosynthetic genes. *AT3G12130*, highly expressed in roots, belongs to the CCCH-type (C3H) zinc finger family, whose members have been implicated in the regulation of secondary metabolism and stress-responsive pathways. Given the central role of cytochrome P450 monooxygenases in triterpenoid oxidation, *AT3G12130* may influence ginsenoside biosynthesis by modulating the expression of downstream P450 genes. Cytochrome P450 enzymes are well known for their involvement in terpenoid, flavonoid, and saponin oxidation [36], and representative members such as *CYP76C3* and *CYP71B31* have been shown to act on triterpenoid substrates in coordination with terpene synthases [37]. Although direct evidence for *AT3G12130* remains lacking, its tissue-specific expression pattern and functional classification support a potential regulatory role in ginsenoside modification.

*SPL9*, another root-enriched transcription factor, is known to regulate auxin signaling by repressing *SAUR19* and influencing cell elongation [38]. Auxin-responsive transcription factors such as bHLH and AUX/IAA have been implicated in ginsenoside biosynthesis [39]. SPL9 is also responsive to BR signaling and can form regulatory complexes with *BZR1*[40], consistent with the upregulation of BR biosynthesis-related genes observed in roots. These findings suggest that *SPL9* may act as a hormone-responsive integrator coordinating developmental and metabolic signals during ginsenoside biosynthesis.

In flowers, *MYB33* and *SPL1* were identified as potential regulators. Although direct links between *MYB33* and ginsenoside biosynthesis have not been reported, *MYB* transcription factors are widely involved in secondary metabolism. In *P. notoginseng*, *PnMYB4* negatively regulates ginsenoside biosynthetic genes[41], and *MYB* overexpression in other species has been shown to enhance triterpenoid production by activating upstream pathways. *SPL1* has been associated with glucose metabolism and energy regulation[42]. Given that glycosylation is a key step in saponin maturation and depends on UDP-sugars and UDP-glycosyltransferases (UGTs)[43], *SPL1* may influence ginsenoside glycosylation patterns in flowers by regulating sugar metabolism or *UGT* expression.

Overall, this study clarifies the spatial and developmental distribution of ginsenosides in *P. notoginseng*, identifies organ- and age-specific regulatory genes, and proposes several core transcription factors potentially involved in ginsenoside biosynthetic regulation. These findings advance the understanding of transcriptional control underlying ginsenoside biosynthesis and provide valuable molecular targets for breeding, metabolic engineering, and the precise utilization of *P. notoginseng* in the food and health industries.

## Conclusions

By integrating targeted metabolomics with multi-organ transcriptomic analyses across developmental stages, this study delineated the spatiotemporal patterns of ginsenoside accumulation in *Panax notoginseng*. Roots and flowers were identified as major sites for the accumulation of distinct ginsenoside types, accompanied by pronounced tissue- and age-specific transcriptional reprogramming. Through expression clustering, transcription factor enrichment, and promoter motif analyses, four transcription factors—*AT3G12130*, *SPL9*, *MYB33*, and *SPL1*—were identified as core candidates associated with tissue-specific ginsenoside biosynthesis. These transcription factors are predicted to regulate key biosynthetic steps, including triterpenoid oxidation and glycosylation, by targeting enzymes such as *CYP716A53v2* and *CYP716A47*. The enrichment of rare ginsenosides, including CK, Rg3, and Rh2, further highlights the potential for tissue-targeted utilization in functional food and nutraceutical applications. Although the proposed regulatory relationships are primarily based on transcriptomic and motif-based evidence, this study provides a systematic framework for understanding the spatial regulation of ginsenoside biosynthesis in *P. notoginseng* and establishes a foundation for future functional validation and metabolic engineering efforts.

## Materials and methods

### Plant Materials

A total of 33 samples, including fresh and healthy roots, stems, leaves, and flowers from one-, two-, and three-year-old plants of *P. notoginseng* were harvested from Wenshan County, Yunnan Province, China, on August 12, 2015. Three biological replicates were collected for each sample type. After harvesting, tissues were immediately flash-frozen in liquid nitrogen and stored at −80 °C for subsequent RNA-seq analysis. For ginsenoside content measurement, approximately 5 g of fresh material was collected, enzyme-inactivated by microwave treatment for 2 minutes, and air-dried at room temperature.

### Determination of ginsenoside contents

Ginsenoside contents in different tissues were quantified using high-performance liquid chromatography (HPLC). An Agilent 1100 HPLC system equipped with a ZORBAX SB-C18 column (4.6 × 250 mm, 5 μm) was used to determine the contents of major ginsenosides, including Rb1, Rb2, Rd, Re, Rg1, and R1. All analyses were performed at 30 °C.

Briefly, approximately 1.00 g of dried tissue powder was weighed and wetted with 90 mL of 70% methanol. After ultrasonic extraction, the volume was adjusted to 100 mL with 70% methanol. The extract was filtered through a 0.45 μm organic membrane prior to injection. A 10 μL sample was loaded onto the column, and elution was performed using a gradient of 0.5% formic acid in water (solvent A) and acetonitrile (solvent B) :8% B for 5 min, 25% B for 23 min, then returning to 8% B for 25 min. The flow rate was 1.0 mL/min, and detection was conducted at 280 nm. Ginsenosides were identified by comparing retention times with authenticated standards.

### RNA-Seq library preparation

Total RNA was extracted from each tissue individually using a modified CTAB method [44]. RNase-free DNase I (Takara) was used to remove residual DNA. RNA quality and integrity were assessed using a NanoDrop-1000 UV-VIS spectrophotometer (NanoDrop) and an Agilent 2100 Bioanalyzer (Agilent Technologies, Palo Alto, CA, USA), respectively. For each RNA sample, at least 20 µg of total RNA at a concentration of ≥ 400 ng/µl was used for cDNA library construction following the manufacturer’s instructions (Illumina, USA). Libraries were constructed and sequenced using the Illumina HiSeq2000 platform.

### RNA-Seq data analysis

Raw reads were preprocessed using Trim_Galore v0.4.4 to remove adapters and low-quality bases [45] Clean reads were aligned to the *P. notoginseng* reference genome using HISAT2 v2.1.0 [46], and unmapped reads were removed with SAMtools v1.8 [47]. Gene-level read counts were calculated using HTSeq v2.0.3 [48]. Differentially expressed genes (DEGs) were identified using DESeq2 v2.13 [49], with an adjusted *p*-value < 0.05 and |log2FoldChange| > 1 as significance thresholds. Expression levels were quantified as fragments per kilobase of transcript per million mapped reads (FPKM) using StringTie v1.3.6 [50]. Genes with CPM > 1 in at least one sample were retained for downstream analysis.

Gene Ontology (GO) terms were assigned using PANNZER2 [51]. GO enrichment was performed with the R package topGO v2.40.0 [52], and terms with *p* < 0.05 were considered significant. To identify cis-regulatory elements (CREs), motif enrichment analysis was performed using HOMER’s findMotifsGenome.pl function with default parameters; motifs with *q* < 0.05 were considered enriched [53]. Transcription factor family classification was based on PlantTFDB annotations via protein sequence similarity [54]. FIMO was used to locate TF-specific binding motifs within the 2 kb upstream regions of candidate genes (*p* < 0.0005) [55]. Functional annotation of ginsenoside biosynthetic enzymes was based on published literature.

## Acknowledgments

This work was supported by Natural Science Foundation of China (U1902205) and a startup grant of Hainan University (RZ2100006631) (to Gao LZ).

## Author contributions

L.G. conceived and supervised the research. F.W., X. G. and Z.C. provided resources and collected the data. W.L. and Y.W. performed the data analysis, interpreted the results, and drafted the manuscript. L.G. revised the manuscript. All authors reviewed the results and approved the final version of the manuscript.

## Data availability

The RNA-seq data analyzed in this study were generated previously by the corresponding author and are publicly available in the National Genomics Data Centre (NGDC) under accession number PRJCA002506. All other relevant data are included in this article and its supplementary materials.

## Supplementary material

Supplementary material is available for this paper. It includes supplementary figures (Figures S1–S4), supplementary tables (Tables S1–S2), and additional methodological details supporting the results and conclusions presented in the main text.

Table S1. Summary of RNA-seq read alignment statistics.

Table S2. Functional annotation of putative target genes regulated by core transcription factors.

---

**Figure S1.**
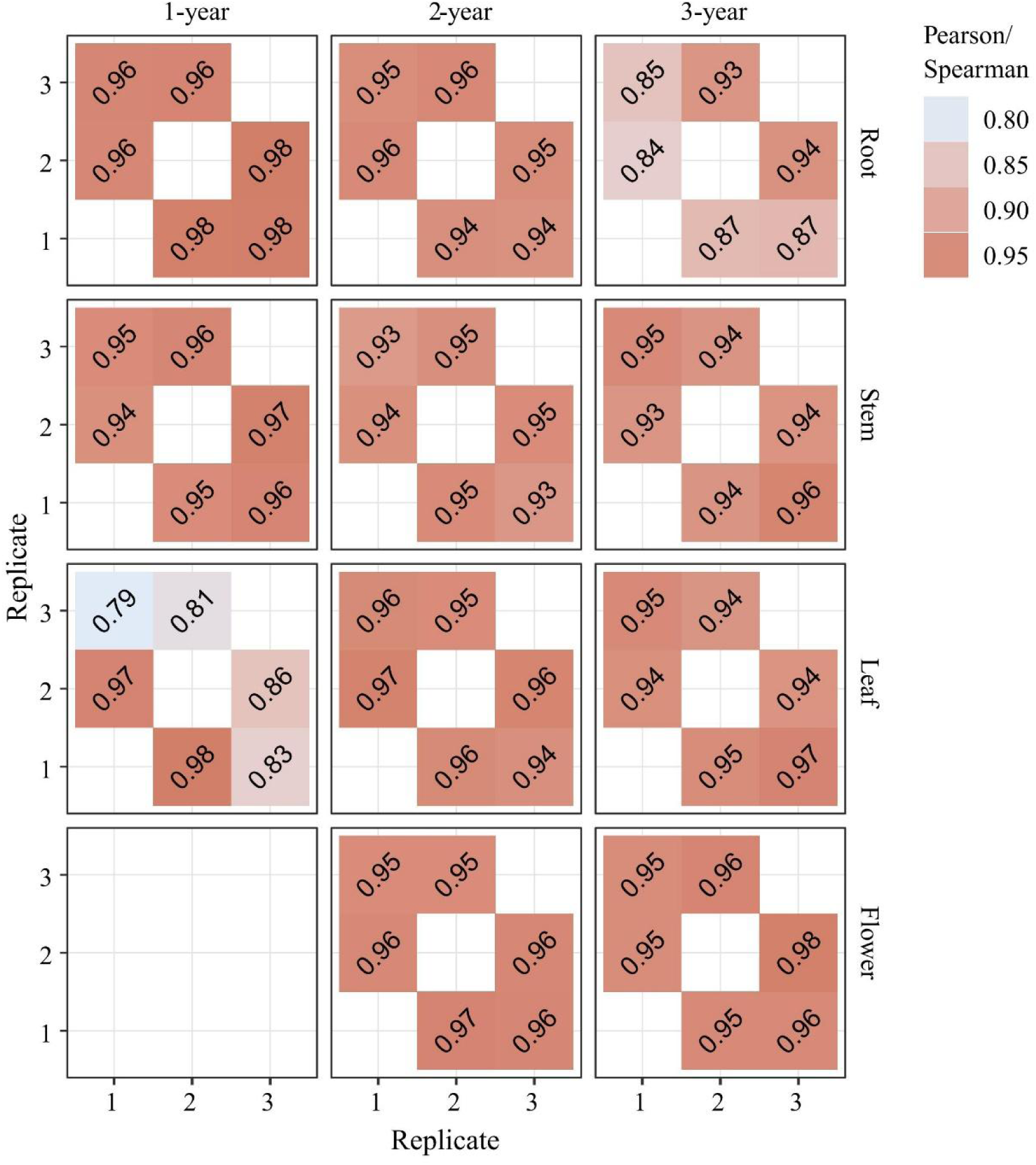
Reproducibility of RNA-seq data among biological replicates. Pairwise correlation analysis was performed to assess the consistency among the three biological replicates for each RNA-seq sample across different tissues and growth years. The upper triangle of the correlation matrix shows Pearson correlation coefficients, while the lower triangle displays Spearman rank correlation coefficients. Color intensity reflects the strength of correlation, with higher values indicating greater reproducibility among replicates.

**Figure S2.**
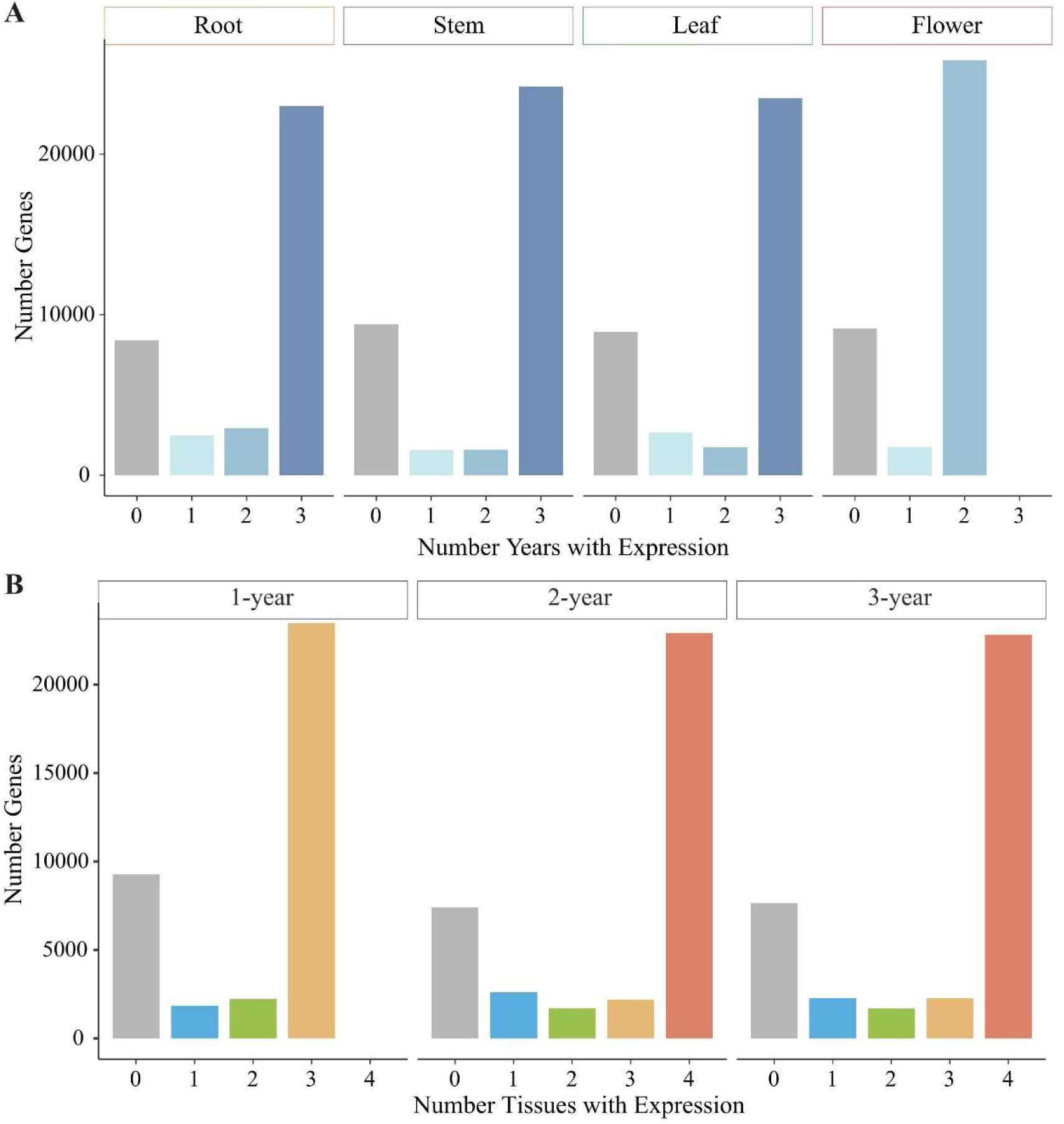
Distribution of expressed genes across tissues and years in *P. notoginseng*. (A) Distribution of expressed genes (CPM > 1) across different tissues within the same growth year. The x-axis indicates the number of tissues in which each gene was detected as expressed. (B) Distribution of expressed genes (CPM > 1) across different growth years within the same tissue. The x-axis indicates the number of years in which each gene was detected as expressed.

**Figure S3.**
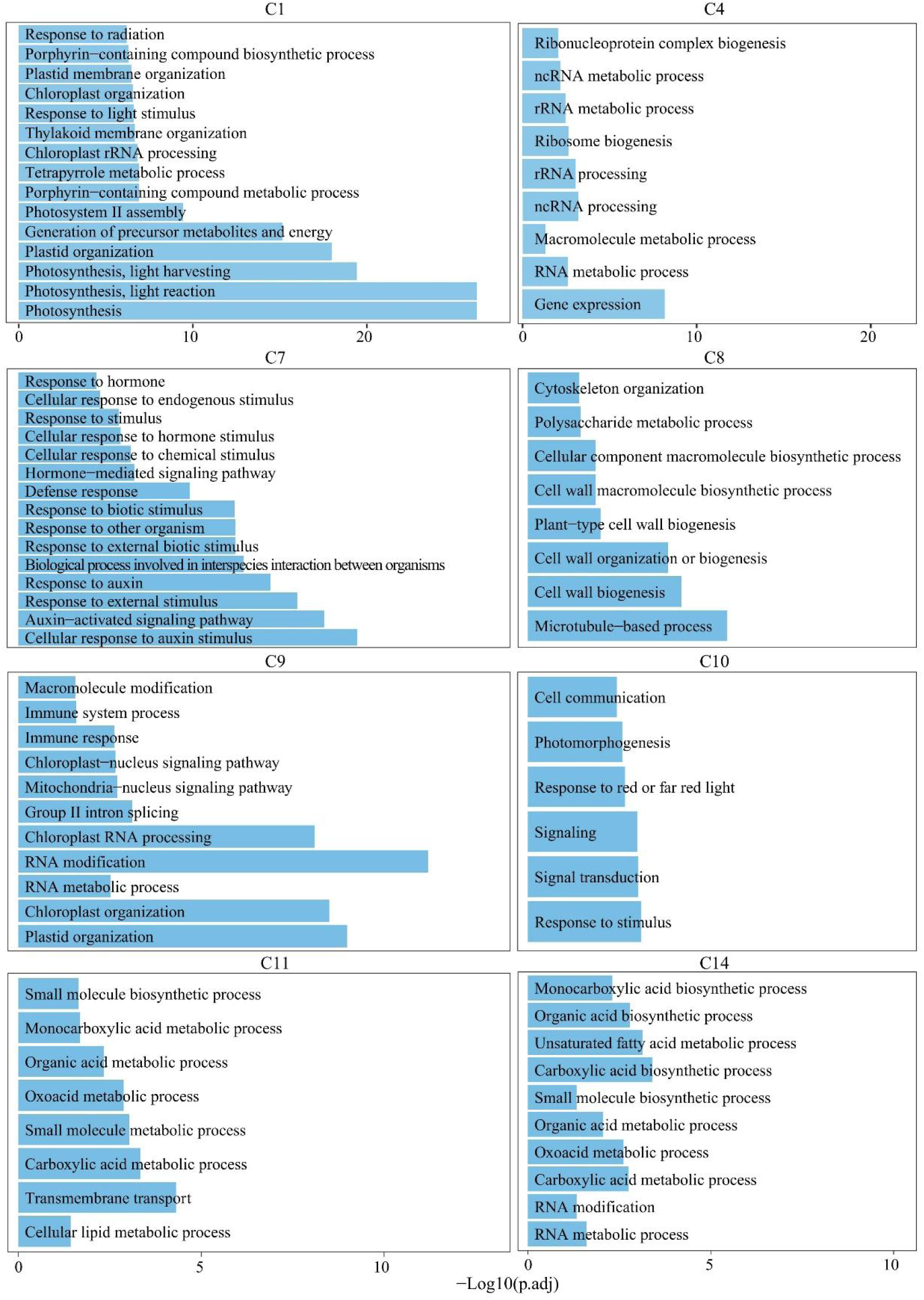
Gene Ontology (GO) enrichment analysis of genes in different expression clusters. Representative enriched biological process terms are shown for each cluster. The color scale indicates enrichment significance, expressed as −log10(q-value) (or enrichment score, as indicated).

**Figure S4.**
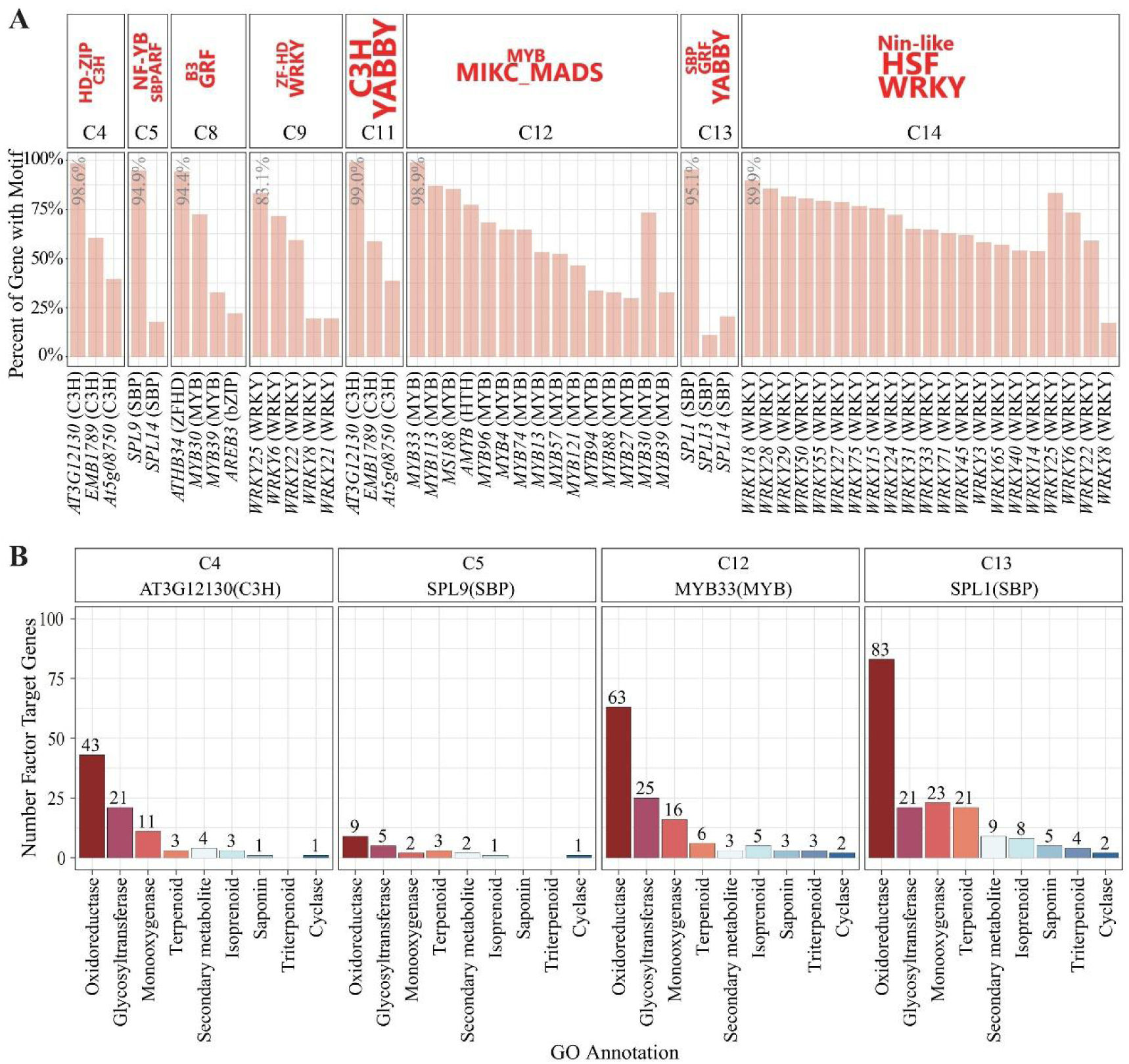
Enrichment analysis of transcription factor binding motifs across gene expression clusters (C1–C15). (A) Identification of significantly enriched transcription factor binding motifs within the 2 kb upstream promoter regions (relative to the transcription start site, TSS) of genes in each cluster. The x-axis indicates the proportion of genes within a given cluster containing the corresponding motif. Transcription factor families that are significantly overrepresented in each cluster are highlighted in red. (B) Gene Ontology (GO) functional annotation of putative target genes regulated by core transcription factors identified in each cluster. Major functional categories are summarized, and detailed GO terms are provided in Table S2.

